# Microbial Reduction of Geogenic and Synthetic Goethite and Hematite

**DOI:** 10.1101/2023.12.05.570092

**Authors:** Edward J. O’Loughlin

## Abstract

The microbial reduction of Fe(III) is a major component of Fe cycling in terrestrial and aquatic environments and is affected by the Fe(III) mineralogy of the system. The majority of the research examining the bioreduction of Fe(III) oxides by Fe(III)-reducing bacteria (IRB) has focused on the reduction of poorly crystalline Fe(III) phases, primarily ferrihydrite; however, crystalline Fe(III) oxides like goethite (α-FeOOH) and hematite (α-Fe_2_O_3_) comprise the majority of Fe(III) oxides in soils. This study examined the bioreduction of goethite and hematite of geogenic and synthetic origin by *Shewanella putrefaciens* CN2, a well-studied model IRB, in laboratory incubations. Overall, the rate and extent of Fe(II) production were greater for goethite than for hematite, and for geogenic Fe(III) oxides relative to their synthetic analogs. Although there was substantial production of Fe(II) (i.e., > 5 mM Fe(II)) in many of the systems, X-ray diffraction analysis of the solids at the end of the incubation did not indicate the formation of any Fe(II)-bearing secondary minerals (e.g., magnetite, siderite, green rust, etc.). The results of this study demonstrate the variability in the extent of bioreduction of geogenic goethite and hematite, and furthermore, that synthetic goethite and hematite may not be good analogs for the biogeochemical behavior of Fe(III) oxides in aquatic and terrestrial environments.

## 1. Introduction

Microbial Fe(III) reduction is a key component of the biogeochemical cycling of Fe in aquatic and terrestrial environments [1-4]. Many forms of Fe(III) can be used as terminal electron acceptors for anaerobic respiration by dissimilatory iron-reducing bacteria (DIRB, which are a phylogenetically diverse group of microorganisms [5-18]) including soluble Fe(III) complexes; structural Fe(III) in aluminosilicate minerals; and Fe(III) oxides, hydroxides, and oxyhydroxides (hereafter collectively referred to as Fe(III) oxides) including akaganeite (β-FeOOH), feroxyhyte (d′-FeOOH), ferrihydrite, goethite (α-FeOOH), hematite (α-Fe_2_O_3_), lepidocrocite (γ-FeOOH), maghemite (γ-Fe_2_O_3_), and ferric green rust [19-34]. The bioreduction of Fe(III) by DIRB can result in the production of a broad range of Fe(II) species including soluble and adsorbed Fe(II), and mineral phases containing structural Fe(II) (e.g., siderite (FeCO_3_), chukanovite [Fe_2_(OH)_2_CO_3_], magnetite (Fe_3_O_4_), green rust, and vivianite [Fe_3_(PO_4_)_2_•8H_2_0], and Fe(II)-bearing clays [10,19,25,26,34-40].

The majority of the research examining the reduction of Fe(III) oxides by DIRB has focused on the reduction of poorly crystalline Fe(III) phases, primarily ferrihydrite. Although crystalline Fe(III) oxides like goethite and hematite comprise the majority of Fe(III) oxides in soils [41], the bioreduction of these phases by DIRB has been less studied. This paper focuses on the microbial reduction of goethite and hematite of geogenic and synthetic origin by *Shewanella putrefaciens* CN2, a well-studied model DIRB originally isolated from subsurface sediment [22].

## 2. Materials and Methods

### 2.1. Geogenic and Synthetic Goethite and Hematite

Synthetic goethite (Bayferrox 910, Lot 3011225) and hematite (Bayferrox 130, Lot 3011225) were obtained from LANXESS Corp., Cologne, Germany. An additional synthetic hematite was obtained from Rockwood Pigments, Inc., Beltsville, MD, USA. Natural Sienna (NS: country of origin, France), Natural Umber (NU: country of origin, France), Natural Red (NR: country of origin, India), Red Ochre (RO: country of origin, France), and Natural Yellow (NY: country of origin, India) were purchased from The Earth Pigments Company, Cortaro, AZ, USA. Dark Ochre (DO: country of origin, Germany) and French Ochre JALS (FOJ: country of origin, France) were purchased from Kremer Pigments, Inc., New York, NY, USA. The mineralogy of the synthetic and geogenic Fe(III) oxides was determined by powder X-ray diffraction (pXRD) with a Rigaku MiniFlex X-ray diffractometer using Ni-filtered Cu Kα radiation, scanned between 5° and 80° 2θ at a speed of 0.1° 2θ min^-1^. The XRD patterns were analyzed with the JADE 9 software package (MDI, Livermore, CA, USA).

### 2.2. Bioreduction Experiments

The bioreduction experiments were conducted as described by O’Loughlin et al. [34]. Briefly, 100 mL of sterile defined mineral medium (DMM) containing 80 mM Fe(III) in the form of one of the geogenic or synthetic Fe(III) oxides, 75 mM formate, 100 µM phosphate, and 100 µM 9,10-anthraquinone-2,6-disulfonate (AQDS) as an electron shuttle in the AQDS-amended systems, was placed in 160-mL serum bottles. The bottles were sealed with rubber septa and aluminum crimp caps and made anoxic by sparging with sterile argon. All experimental systems were prepared in duplicate. The bottles were inoculated with *S. putrefaciens* CN32 (American Type Culture Collection BAA-543) (prepared as described in O’Loughlin et al. [42]) at a density of ∼5 × 10^9^ cells mL^-1^ and placed on a roller drum and incubated at 30 °C in the dark. Samples of the suspensions for monitoring Fe(II) production and identification of secondary minerals by pXRD were collected with sterile syringes and analyzed as described by O’Loughlin et al. [34]. Briefly, samples for Fe(II) analysis were prepared by adding 0.75 mL of anoxic 1 M HCl to 0.25 mL of suspension and the Fe(II) concentration was determined by the ferrozine assay [43]. Samples for pXRD analysis were collected by filtration on 25-mm, 0.22-µm nylon filters and covered with 8.4-µm-thick Kapton^®^ film under anoxic conditions.

## 3. Results and Discussion

### 3.1. Characterization of Geogenic and Synthetic Goethite and Hematite

All three of the synthetic Fe(III) oxides are highly crystalline and show no indication of crystalline constituents other than the phase indicated by the manufacturer; i.e., goethite in the case of Bayferrox 910 and hematite for Bayferrox 310 and Rockwood hematite (Figure 1).

**Figure 1.**
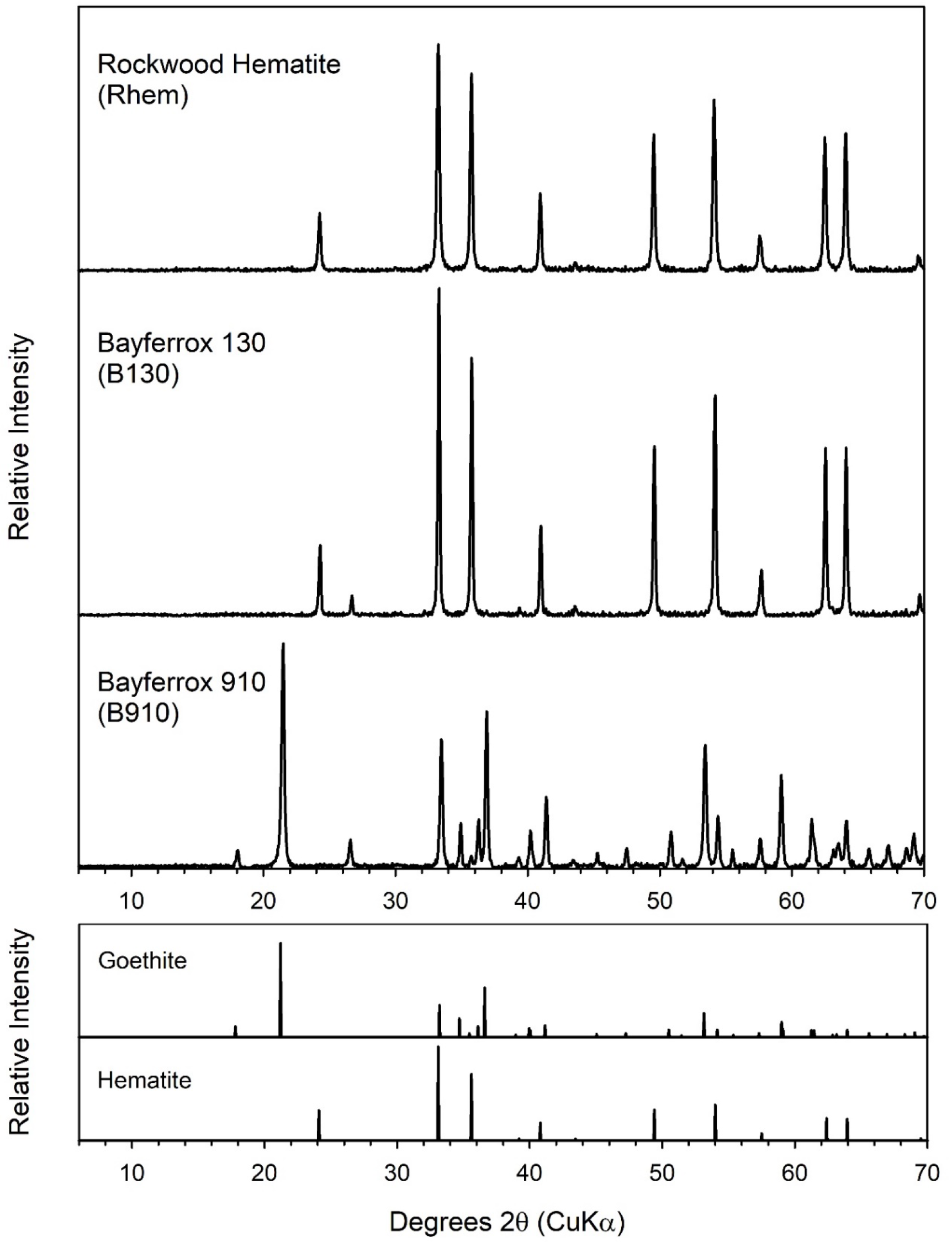
pXRD analysis of the synthetic goethite and hematites used in this study.

Geogenic iron oxides have been used as pigments since prehistoric times and are commonly classified as ochres, siennas, umbers, and blacks [41,44,45]. In contrast to the synthetic phases, the geogenic iron oxides used in this study consist of mixtures of several minerals. XRD analysis of the geogenic pigments (Figures 2 and 3) indicated that the Fe(III) oxide component was goethite (NS, NY, NU, DO, and FOJ), hematite (NR), or a mixture of both (RO). It is important to note however, that pXRD is most sensitive to crystalline materials, therefore, the presence of poorly crystalline and nano-scale Fe(III) oxide phases cannot be excluded. Other crystalline components included quartz (NS, NY, NU, DO, FOJ, and RO) and kaolinite (NS, NR, DO, FOJ, and RO) (Table 1), consistent with previous characterization of similar ochres, siennas, and umbers [46-49].

**Table 1.**
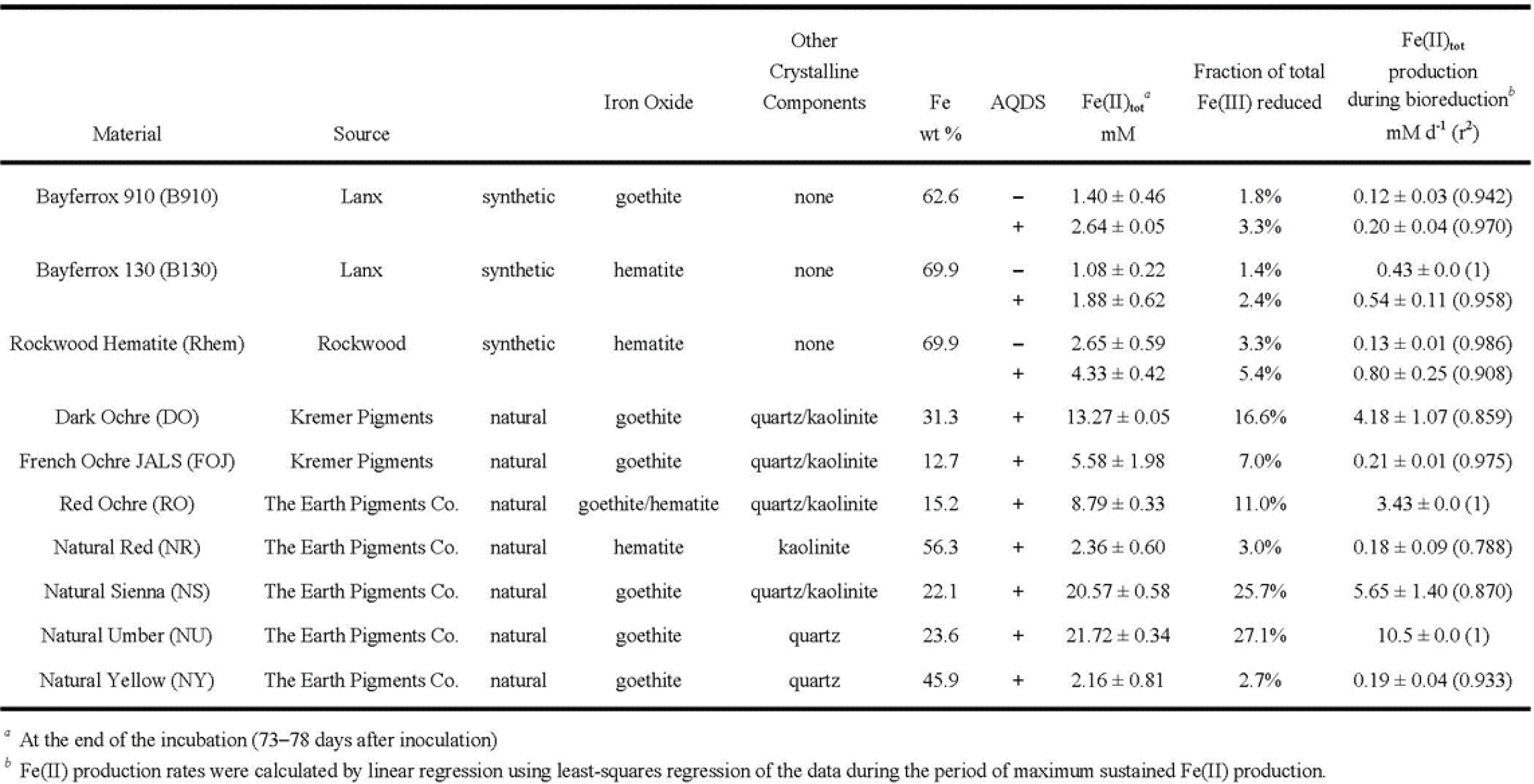
Fe(III) oxide source and mineralogy, Fe content, Fe(II) production, fraction Fe(III) reduced, and maximum Fe(II) production rates.

**Figure 2.**
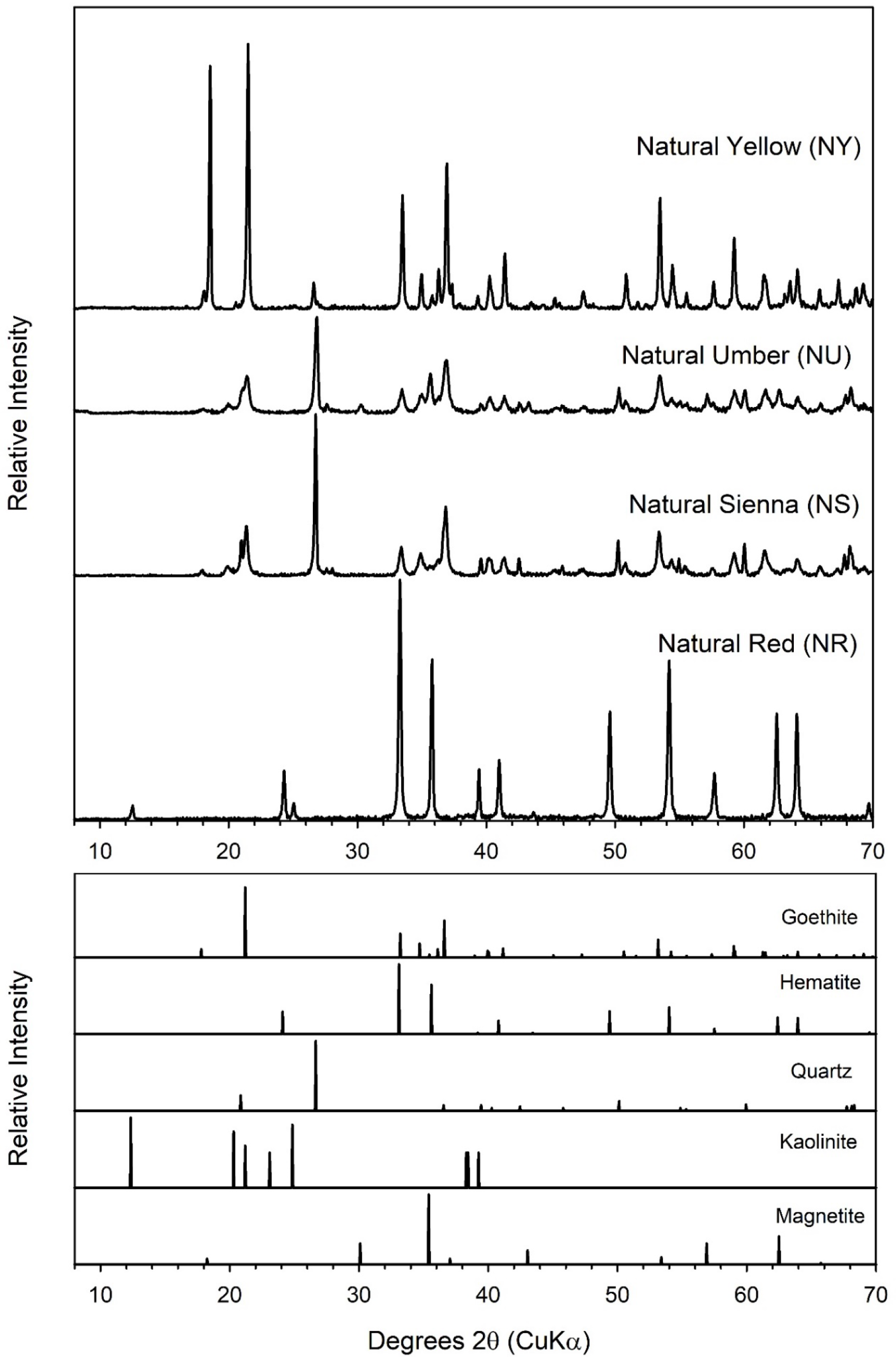
pXRD analysis of the geogenic iron oxides used in this study.

**Figure 3.**
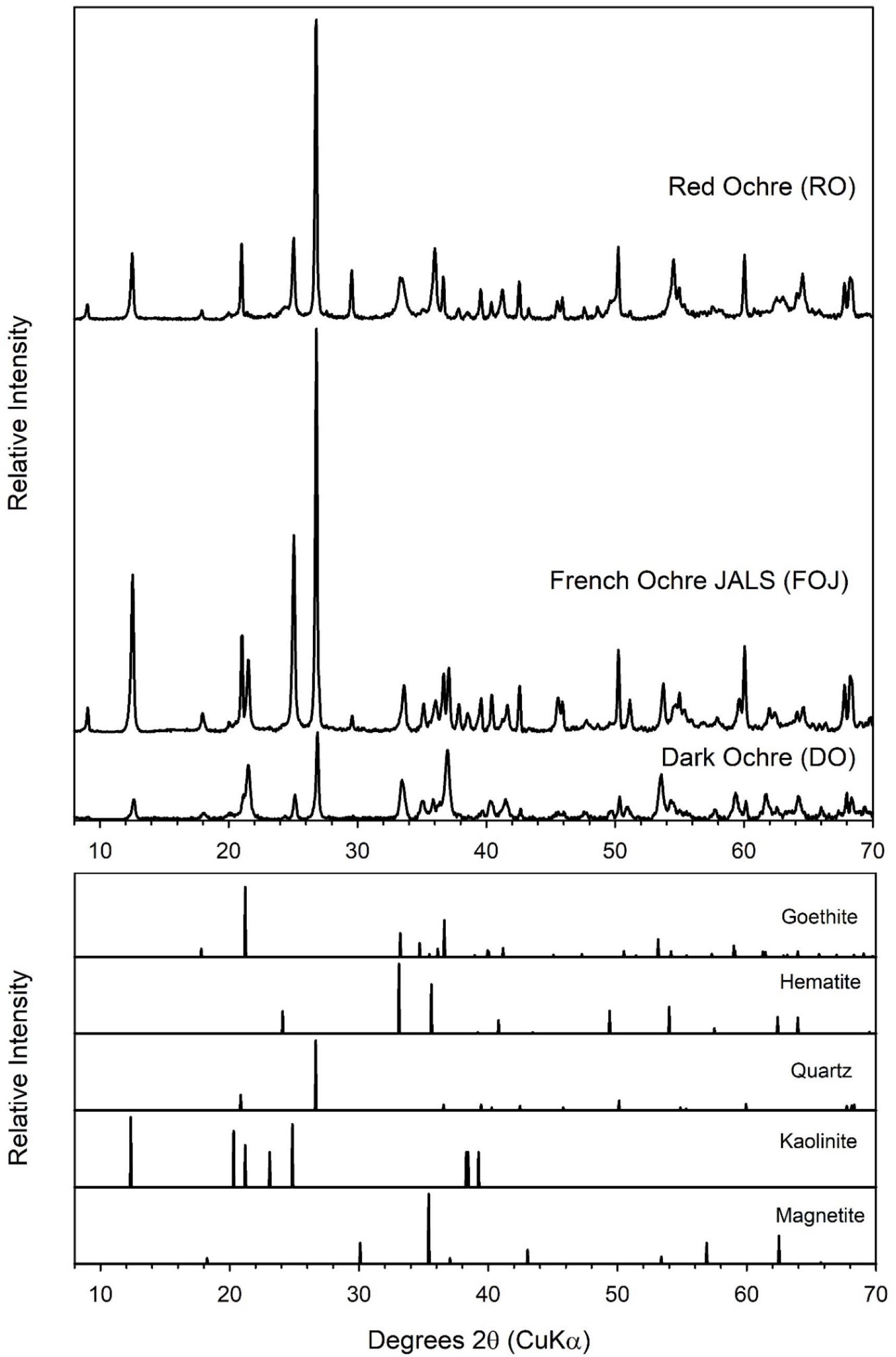
pXRD analysis of the ochres used in this study.

### 3.2. Bioreduction of Geogenic and Synthetic Goethite and Hematite

The extent of Fe(III) reduction varied greatly among the systems containing goethite (Figure 4 and Table 1). The bioavailability of Fe(III) in the synthetic goethite B910 was extremely low, with only 3.3% of the Fe(III) reduced to Fe(II) over a period of 78 days. This is substantially lower than the extent of Fe(III) reduction reported by O’Loughlin et al. [34] for a different synthetic goethite under the same experimental conditions (13% within 78 days). Although they are both synthetic phases, the commercial goethite B910 is highly crystalline (Figure 1) while the lab-synthesized goethite prepared by O’Loughlin et al. [34] is poorly crystalline in comparison (Figure 7 in [34]). It is likely that the differences in the reducibility of Fe(III) in the two synthetic goethites is due to differences in their crystallinity, as the bioavailability of Fe(III) in Fe(III) oxides tends to decrease as the crystallinity of the phase increases. This general trend is also apparent among the geogenic goethite systems. Slower rates and lower overall extents of Fe(II) production were observed during the bioreduction NY and FOJ, which have comparatively higher goethite crystallinity, and the systems with goethites of lower crystallinity (DO, FS, and NU) had faster rates and greater extents of Fe(II) production (Figures 2, 3, and 4 and Table 1).

**Figure 4.**
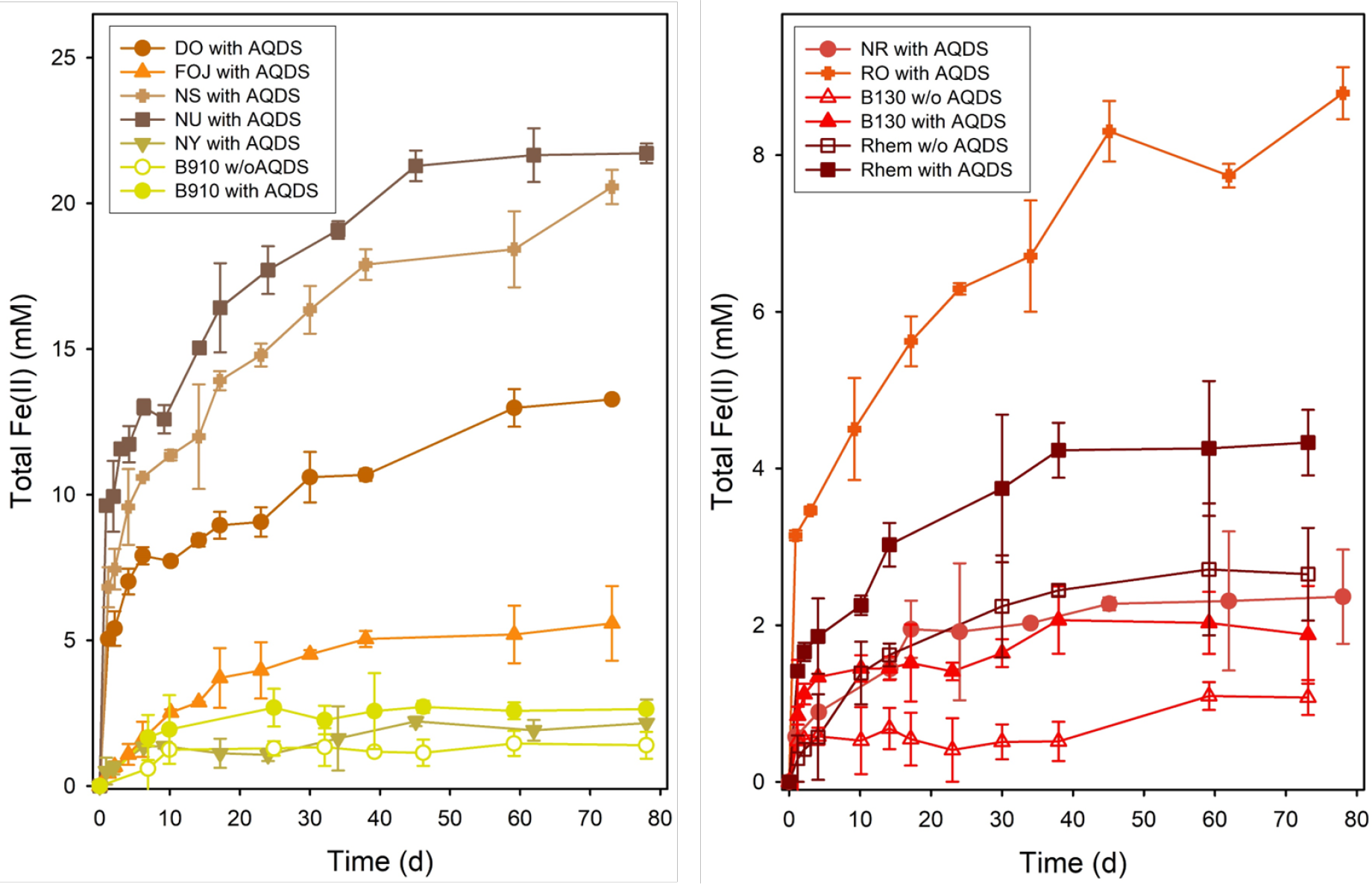
Fe(II) production during the bioreduction in the systems containing synthetic or geogenic goethite (left) or hematite (right).

In the systems containing only hematite, there was little difference between the rates and extents of reduction among the geogenic and synthetic materials (Figure 4 and Table 1), with 2.4% reduction of B130 (synthetic), 3.0% reduction of NR (geogenic), and 5.4% reduction of Rhem (synthetic). Overall, the bioavailability of Fe(III) in the hematite samples was lower than in the goethite samples, consistent with previous studies showing hematite to be generally less susceptible to microbial reduction than goethite [22,28,31,34,50,51]. Compared to the other hematite-containing systems, a faster rate and greater extent of Fe(II) production was observed with RO (Figure 4 and Table 1); however this is likely due to the fact that this material contained both hematite and goethite, with the Fe(III) in the goethite component likely being more reducible.

The presence of electron shuttles (soluble compounds or materials that can be reversibly oxidized and reduced) has been shown to enhance the rate, and often the extent, of the bioreduction of a wide range of Fe(III) oxides [10,22,31,52-60] and AQDS is often used as a model electron shuttle as it is seen as an analog for the quinone groups in natural organic matter [61]. A comparison of the bioreduction of the synthetic goethite and hematites in the presence and absence of AQDS indicates a substantial increase in both the rate and extent of Fe(II) production with the addition of AQDS (Table 1), consistent with previous studies.

### 3.3. Secondary Minerals

During the microbial reduction of Fe(III) oxides, the production of Fe(II) is often accompanied by the formation of Fe(II)-bearing secondary minerals, which can include magnetite, siderite, vivianite, green rust, and chukanovite. Previous studies of the bioreduction of goethite have reported the formation of siderite, vivianite, and chukanovite [22,34,37,62-64], and siderite, magnetite, and vivianite have been reported as products of the bioreduction of hematite [34,35,37,65,66]. However, pXRD analysis of the solids in the bioreduction systems at the end of the incubations did not indicate the formation of any Fe(II) secondary mineral in any system (Figures 6 and 7). Since XRD is sensitive to the crystallinity of solid phases, the formation of poorly crystalline or nanoscale secondary minerals cannot be ruled out.

**Figure 5.**
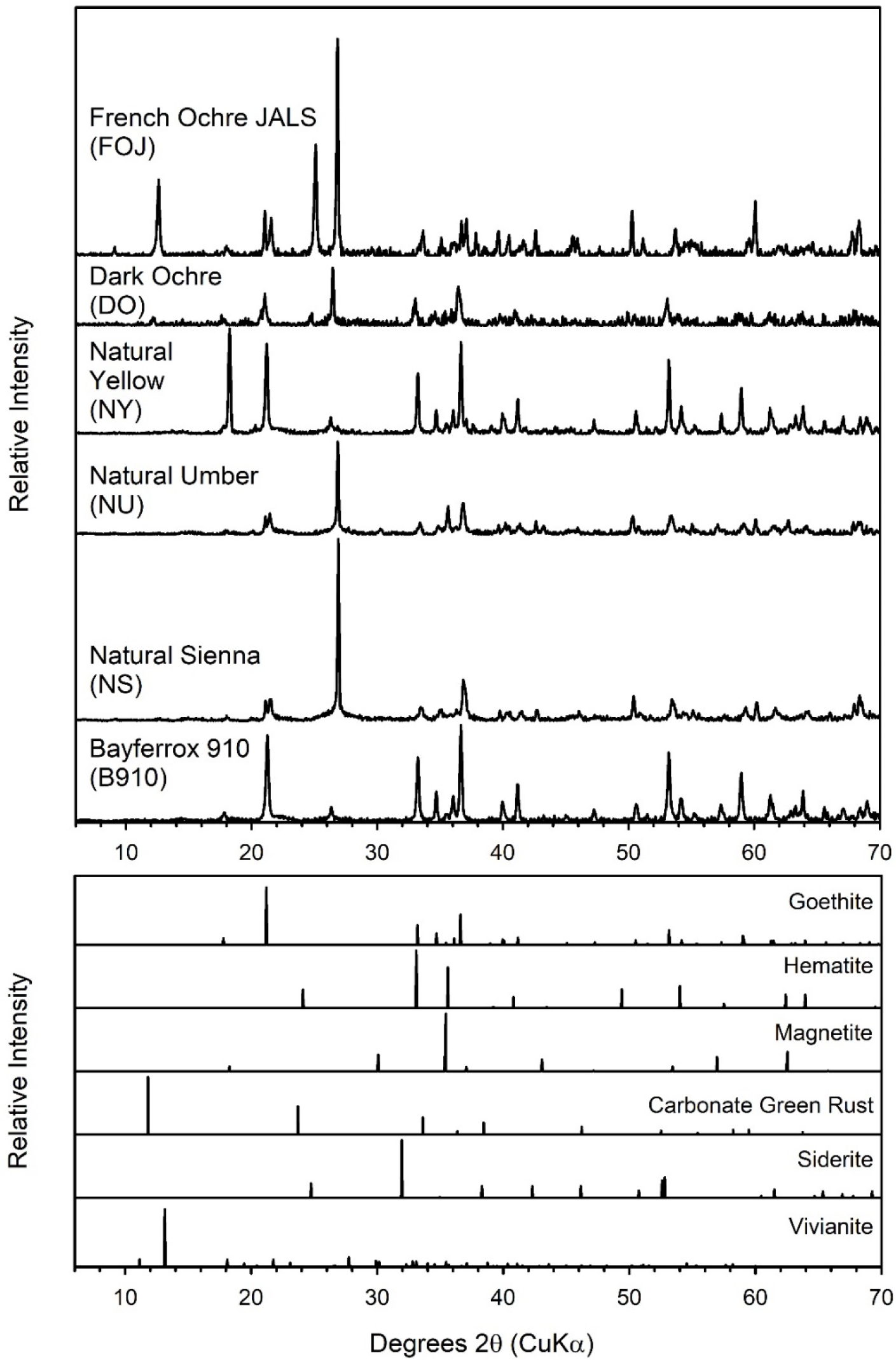
pXRD analysis of the solids in the geogenic and synthetic goethite bioreduction systems at the end of the incubations.

**Figure 6.**
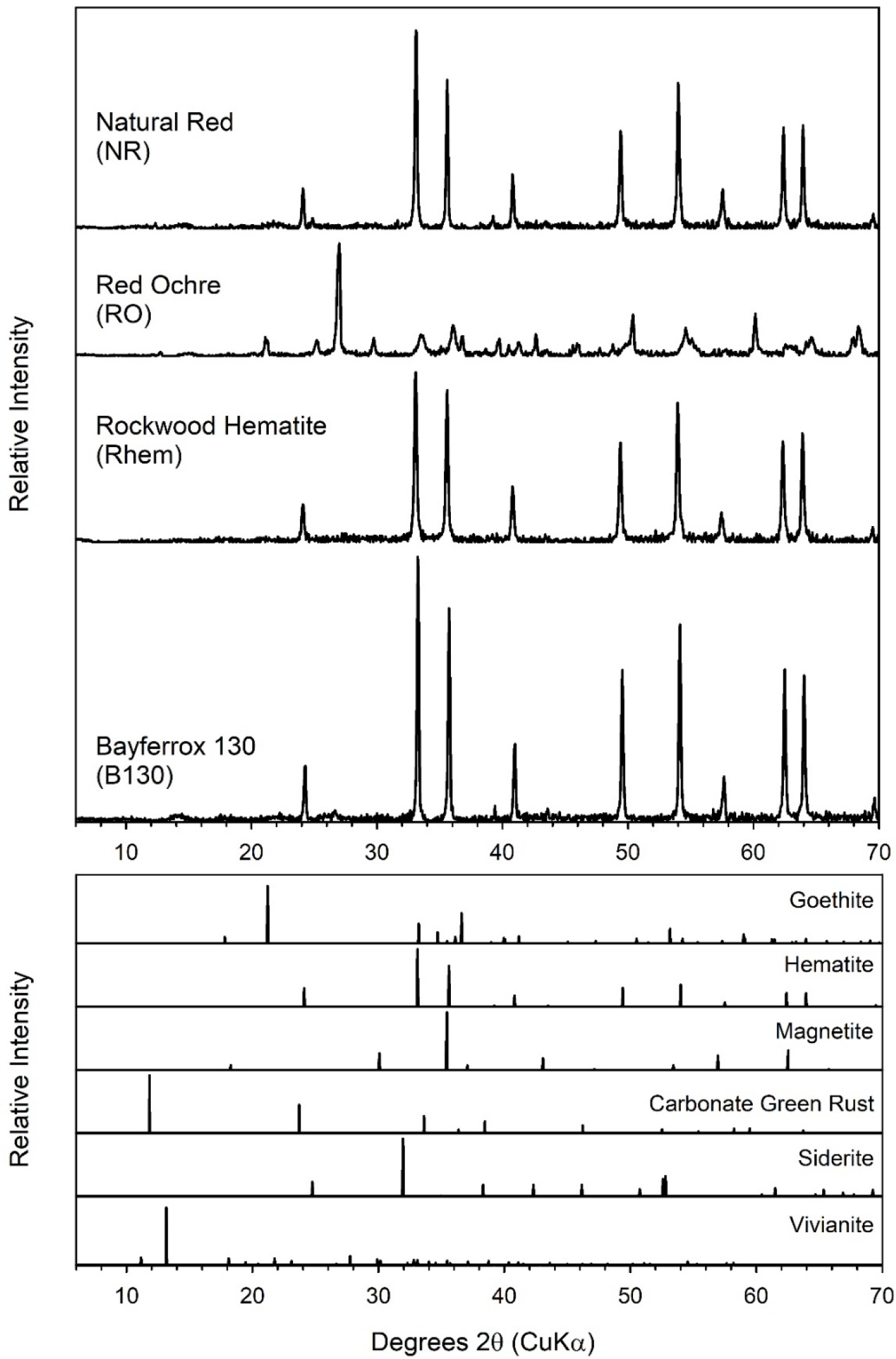
pXRD analysis of the solids in the geogenic and synthetic hematite bioreduction systems at the end of the incubations.

### 3.4. Conclusions

The results of this study show clear differences in the rates and extent of Fe(II) production during the bioreduction by *S. putrefaciens* CN32 of goethite and hematite of synthetic and geogenic origin. These results are consistent with a previous study by Zachara et al., showing that goethite and hematite from geogenic sources were more readily bioreduced than synthetic analogs, which they attributed to crystalline disorder and microheterogeneities [22]. Geogenic Fe(III) oxides often exhibit significant cation substitution by a wide range of divalent and trivalent metal cations; e.g., geogenic goethites may have up to 36% isomorphic substitution of Al^3+^ for Fe^3+^ on a mole basis [41]. However, cation substitution is not a likely factor in the greater bioreducibility of natural goethites, as substitution of Al^3+^ for Fe^3+^ has been shown to have either no effect or to inhibit microbial reduction of goethite [67-69]. On the other hand, synthetic Fe(III) oxides may contain constituents that can affect their biogeochemical behavior due to the procedures used to control their synthesis and properties. For example, trace levels of phosphate (i.e., 0.2 mass%) in a commercial synthetic lepidocrocite decreased the rate of Fe(II) production during bioreduction, but ultimately lead to a greater extent of Fe(III) reduction and differences in the formation of Fe(II)-bearing secondary minerals [70]. Overall, the comparatively lower bioreducibility of synthetic Fe(III) oxide phases relative to their geogenic analogs, suggests that they may not be representative of the biogeochemical reactivity of Fe(III) oxides in aquatic and terrestrial environments.

©2023. This manuscript has been created by UChicago Argonne, LLC, Operator of Argonne National Laboratory (“Argonne”). Argonne, a U.S. Department of Energy Office of Science laboratory, is operated under Contract No. DE-AC02-06CH11357. The U.S. Government retains for itself, and others acting on its behalf, a paid-up nonexclusive, irrevocable worldwide license in said article to reproduce, prepare derivative works, distribute copies to the public, and perform publicly and display publicly, by or on behalf of the Government. The Department of Energy will provide public access to these results of federally sponsored research in accordance with the DOE Public Access Plan. http://energy.gov/downloads/doe-public-access-plan

